# Distinct behavior of the little finger during the vertical translation of an unsteady thumb platform while grasping

**DOI:** 10.1101/2021.05.28.446096

**Authors:** Banuvathy Rajakumar, SKM Varadhan

**Affiliations:** Department of Applied Mechanics, Indian Institute of Technology Madras, Chennai, India

**Author notes:** **Corresponding Author:** Dr Varadhan SKM, Assistant professor, Department of Applied Mechanics, Indian Institute of Technology Madras, Chennai - 600036, Tamil Nadu.

**Keywords:** Thumb, Peripheral fingers, Grasping, Object stabilization

## Abstract

Object stabilization while grasping is a common topic of research in motor control and robotics. Forces produced by the peripheral fingers (index and little) play a crucial role in sustaining the rotational equilibrium of a handheld object. In this study, we examined the contribution of the peripheral fingers towards object stabilization when the rotational equilibrium is disturbed. For this purpose, the thumb was placed over an unsteady platform and vertically translated. The task was to trace a trapezoid or an inverted trapezoid pattern by moving the thumb platform in the vertical direction. The thumb displacement data served as visual feedback to trace the pattern displayed. Participants were instructed to maintain the handle in static equilibrium at all times. We observed that the change in the normal force of the little finger due to the downward translation of the thumb was significantly greater than the change in the normal force of the index finger due to the upward translation. We speculate that morphological correlations (between thumb and little finger) during the displacement of the thumb might be a reason for such large increases in the little finger forces.

## Introduction

In everyday life, among the various activities performed by the human hand, grasping an object and maintaining the object in static equilibrium is quite common^1^. Force distribution on the handheld object varies systematically depending on the shape^2^, mass^3^, location of the thumb^4^, and surface property^5^ of the grasped object. Previous studies on the static multi-finger prehension found that the index finger produced a greater share of normal force, followed by the middle, ring, and little fingers^6,7^. The contribution of the little finger to maintain the grip strength was comparatively lesser than the other fingers during the static holding of an object^8^.

Several studies have investigated the scaling of normal forces when changes were imparted to the object orientation^9^, object width^10^, friction^11^, and external torque^12^ of an object held with all five fingers (prismatic precision grip). In studies where external torque was introduced to the handle, either a pronation or supination moment was required to counter-balance the rotation caused. The normal forces produced by the radial fingers (index and middle) contributed in producing the pronation moment during clockwise perturbation. In contrast, the normal forces of the ulnar fingers (ring and little) were involved in producing the supination moment during anticlockwise perturbation. According to the mechanical advantage hypothesis (MAH), it was expected that the index and little fingers with the larger moment arms for normal forces would produce greater normal force than the middle and ring fingers during tasks that required pronation and supination moments respectively, to stabilize the object ^13,14^.

From literature, it is known that the role played by the peripheral fingers (index and little) differs from the central fingers (middle and ring) according to the task requirement^12^. The forces generated by the central fingers varied depending on both the load and torque changes to the grasped object^12,15^. In contrast, studies have reported an increase in the normal forces of peripheral fingers for the tasks that required the maintenance of the rotational equilibrium of the handle^16,17^. Hence, the peripheral fingers were given special attention when torque changes were introduced to the handheld object.

In our preliminary study^18^, torque changes were incorporated in the handle by placing the thumb on a slider platform matching the midline between middle and ring fingers (henceforth, called as “HOME” position in the rest of the manuscript). Since the mechanical constraint to fix the slider platform was removed, the tangential force of the thumb dropped. In order to compensate for the drop, a supination moment was required to be produced by the rest of the fingers. Although we expected to see a larger normal force contribution by the little finger, the ulnar fingers exerted a comparable normal force. However, the normal forces produced by the ulnar fingers were greater than the normal forces produced by the radial fingers, as expected.

From these studies, it is evident that the contribution of the fingers varied depending on the objects handled. Certain objects in real life require vertical motion of the thumb for their operation. For example, while aspirating a sample fluid using specific models of pipette controllers, the pipette has to be held in a vertical orientation while making a fine vertical adjustment using the thumb. In such tasks, the participation of peripheral fingertip forces is critical in producing greater normal force to overcome the torque changes due to the thumb translation. This idea of providing freedom to the thumb emerged by observing the working of such objects that have a vertical tuner on the thumb side for their operation. Thus, in the current study, a handle with an unsteady thumb platform was designed to examine the contribution of index and little finger forces in re-establishing the rotational equilibrium due to thumb motion. The task involved moving this thumb platform towards index, and little finger ends.

Some studies have examined the individual fingertip forces at discrete locations of the thumb during the static holding of the handle^19^. To the best of our knowledge, none of these studies have focused on the force distribution of peripheral fingers when the thumb is mounted on an unsteady platform and held at different positions. This study will give a better understanding of the morphological relationship of the peripheral fingers with the thumb to overcome the torque changes caused due to the unsteady thumb platform held at various positions.

When the platform is held steady at the HOME position, we expected to see a higher normal force by the little finger when compared with the index finger (based on our previous study). In this manuscript, we restrict our attention on investigating the change in the normal force experienced by the peripheral fingers due to the upward and downward thumb translations from HOME. Additionally, we also checked the absolute magnitude of the normal force exerted by the index and little fingers at the end of thumb translations from the HOME position.

Due to the vertical translation of the thumb platform towards the index and little finger end from the HOME position, there are four possible ways by which normal forces of the peripheral fingers may vary to maintain the rotational equilibrium. The first possibility is that the absolute normal force of the index and little finger will be equivalent when the thumb is moved to the TOP and BOTTOM positions, respectively. This means that there will be a greater *change* in the normal force of the index finger than that of the little finger (see Fig.1). The second option is that the *change* in the normal force of both fingers will be comparable, which means the absolute normal force will still be greater for the little finger. The third option will be a greater absolute normal force and *change* in the normal force of the little finger when the thumb slider reaches the BOTTOM position than those for the index finger when the slider reaches the TOP position. The final possibility is that the absolute normal force and *change* in the normal force of the index finger will substantially increase as the slider reaches the TOP than that for the little finger when the slider is at the BOTTOM position.

**Figure 1.**
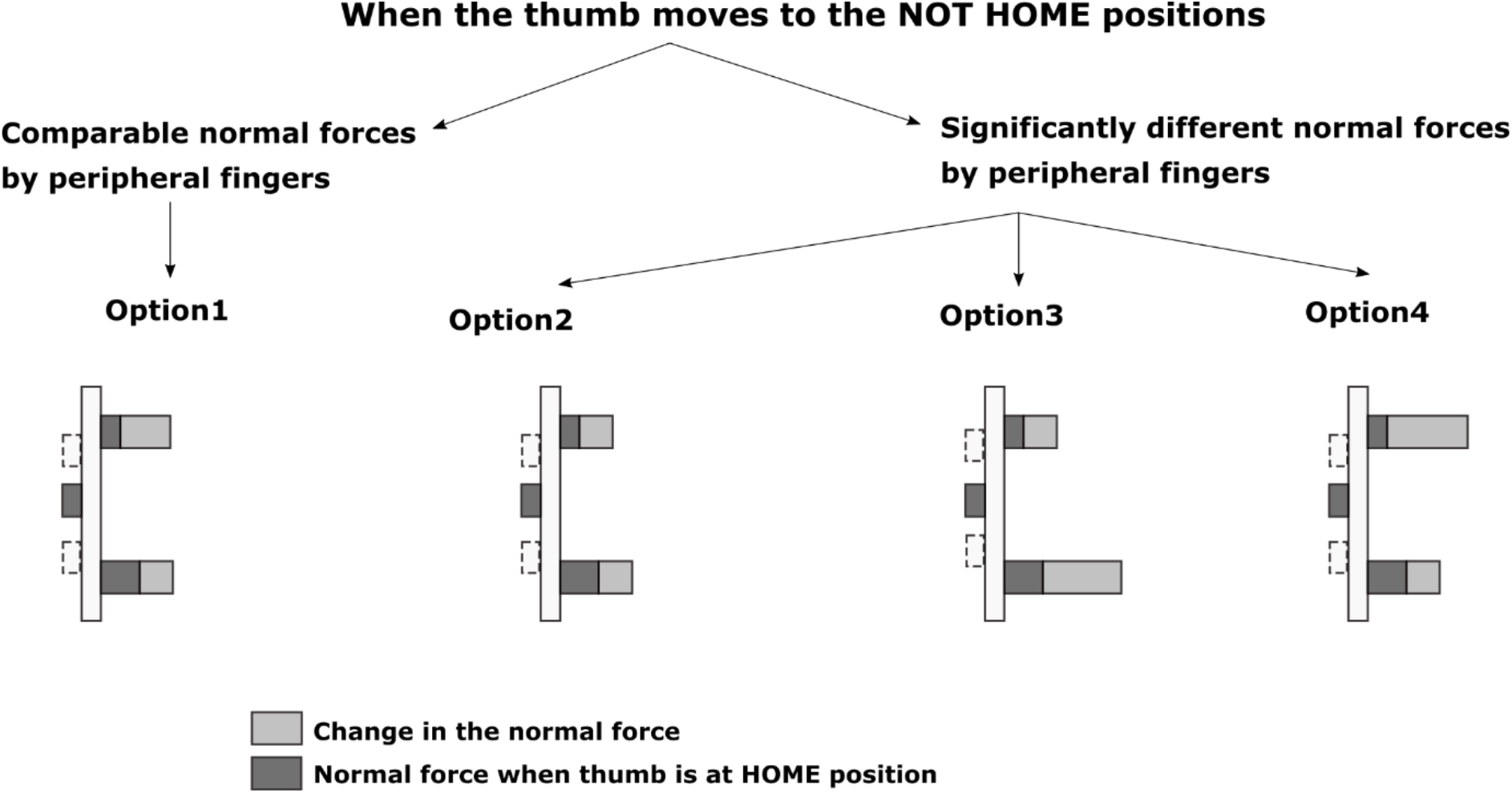
Pictorial representation of four different options shows the normal force pattern of peripheral fingers when the thumb remains at other positions. This figure shows the magnitude of normal force (dark grey) when the thumb was at HOME and the change in the normal force (light grey) experienced by the peripheral fingers when the thumb shifted to TOP & BOTTOM positions. The dashed line represents the TOP & BOTTOM positions of the thumb. In all four options, the absolute normal force exerted by the little finger would be significantly greater than the index finger when the thumb was at the static HOME position. **Option 1:** The *change* in the index finger normal force due to upward translation of the thumb from HOME would be significantly greater than the change in the little finger normal force due to downward translation of the thumb from HOME. Thus, the absolute normal force by the peripheral fingers would be equivalent when the thumb was held at TOP & BOTTOM positions. **Option 2:** The *change* in the normal force of the peripheral fingers remained equivalent. The absolute normal force produced by the index finger (thumb at TOP position) would be significantly lesser than the little finger normal force (when the thumb was at BOTTOM position). **Option 3:** The *change* in the index finger normal force due to upward translation of the thumb from HOME would be significantly lesser than the change in the little finger normal force due to downward translation of the thumb from HOME. Thus, the absolute normal force produced by the index finger when the thumb was at the TOP position would be significantly lesser than the little finger normal force when the thumb was at the BOTTOM position. **Option 4:** The *change* in the index finger normal force due to the upward translation would be significantly greater than the change in the little finger normal force due to the downward translation. Thus, the absolute normal force produced by the index finger (thumb at TOP position) would be significantly greater than the little finger normal force (thumb at BOTTOM position) at the end of translations.

Since the magnitude of thumb displacement in either direction from the HOME position remained the same, we expected that the second option would most likely be true. Hence, we hypothesized that the *change* in the normal forces of the peripheral fingers would be comparable (Hypothesis H1). Furthermore, we expected that the absolute normal force of the little finger produced at the end of the downward translation of the thumb would be greater than the index finger normal force produced at the end of the upward translation of the thumb. Also, we anticipated that there would be an increase in the normal force of the thumb to compensate for the rise in the normal force of the peripheral fingers when the thumb moved away from HOME. For this reason, we hypothesized that the thumb normal force would show a significant increase when the thumb platform reached various positions away from HOME (Hypothesis H2).

## Materials and Methods

### Participants

Twelve right-hand dominant male volunteers (mean ± standard deviation Age: 22.6±2.4 years, Height:173.4±6.4cm, Weight:70.5.3±9.7kg, Hand-length:19±0.6cm, and Hand-width:9.5±0.6cm) participated in this study. Participants did not have any history of hand injuries or neurological disorders.

### Ethical Approval

The experimental procedures were approved by the Institutional ethics committee of Indian Institute of Technology Madras (Approval number: IEC/2018-03/SKM-2/05). Written informed consent was obtained from all participants before the start of the experiment.

### Experimental setup

An instrumented five-finger prehension handle made of aluminum was designed for performing this experiment, shown in Fig.2A. The thumb side of the handle had a vertical railing over which a slider platform was placed. We measured fingertip forces and moments by mounting five six-component force/torque sensors (Model Nano 17, Force resolution: Tangential: 0.0125N, Normal: 0.0125N, ATI Industrial Automation, NC, USA). The sensor for the thumb was mounted on the slider platform, and other sensors were mounted on the side without railing.

**Figure 2.**
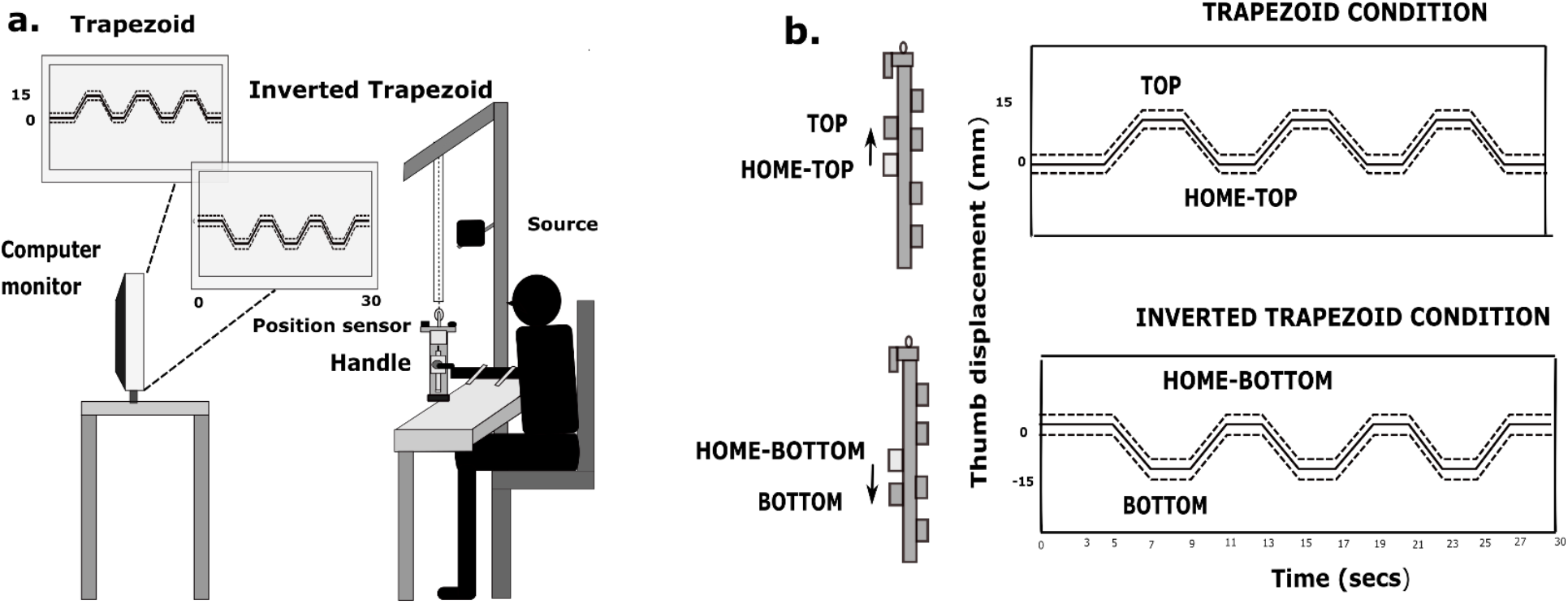
Schematic diagram of the experimental setup with participant holding the five finger prehensile handle with the thumb at four static positions. (A) A computer monitor with the trapezoid and inverted trapezoid patterns was shown to the participant at a distance of 1.5m away. The mass of the handle including the counterweight was 0.535 kg. A rectangular aluminum counterweight of 0.035 kg was placed close to the bottom of the handle on the thumb side to move the whole object’s center of mass close to the geometric center of the handle. The mass of the slider platform alone was 0.101 kg. The friction between the slider and railing was kept minimal by regularly cleaning and lubricating the ball bearing in the slider. In the trial belonging to trapezoid condition, three trapezoids (thick line) were shown consecutively, one after the other. Similarly, in the trial belonging to the inverted trapezoid condition, three inverted trapezoid patterns were shown. Error margins (dashed line) were shown for both the patterns. Complete design of the experimental handle can be referred from the diagram mentioned in the preliminary study (B) Schematic diagram of the handle displaying the position of the thumb at four static positions: TOP, HOME-TOP, HOME-BOTTOM, BOTTOM. The static ‘flat’ portion traced by placing the thumb at the HOME position during trapezoid and inverted trapezoid conditions were called HOME-TOP and HOME-BOTTOM. The above diagram signifies the displacement of the thumb 1.5 cm above, from the HOME-TOP position (no shade) to the TOP position (grey shaded) during trapezoid condition. While in the inverted trapezoid condition, displacement of thumb 1.5 cm below, from the HOME-BOTTOM position (no shade) to the BOTTOM position (grey shaded). A single trial with 30 seconds trial duration on X-axis and 1.5 cm (or 15mm) thumb displacement on the Y-axis for both conditions is shown on the right panel.

A laser displacement sensor (resolution: 5μm; OADM 12U6460, Baumer, India) was attached on top of the handle towards thumb side to measure the vertical displacement of the thumb platform. Towards the participant side, a spirit level with a bull’s eye was provided. On the other side, an electromagnetic tracking sensor (Resolution 1.27 microns, Model: Liberty Standard sensor, Polhemus Inc., USA) was mounted to measure the position and orientation of the handle with reference to the source. The force/torque (thirty channels) and displacement data (single channel) was synchronized with six channels of digital data from the electromagnetic tracker using a customized LabVIEW program. A more complete description of the experimental handle design with the diagram may be found in our earlier publication^18^

### Experimental procedure

Participants were asked to wash their hands with soap and towel-dry before the start of the experiment. They were required to sit comfortably with the forearm resting on the table-top. The right upper arm was abducted approximately 45° in the frontal plane, flexed 45° in the sagittal plane with the elbow flexed approximately 90°. In order to have a natural grasping position, the forearm was supinated to 90°. The movements of the forearm and wrist were restricted by strapping them to the table-top with velcro.

The experiment involved performing a task that consists of two different conditions: tracing trapezoid and inverted trapezoid patterns, as shown in Fig.2B. A template pattern was displayed on the participant’s computer monitor. At 0.5 cm above and below the pattern, dotted lines parallel to the pattern were shown. These dotted lines acted as acceptable error margins. The vertical displacement data of the thumb served as visual-feedback in real-time to trace the pattern displayed on the monitor. For the first five seconds of all the trials, the participants had to position the slider platform steady at the HOME position. This would then be followed by tracing the trapezoid or inverted trapezoid pattern depending on the condition.

In the trapezoid condition, the participants were required to translate the slider platform vertically upwards for 1.5 cm from the HOME position. This involved tracing the “up-ramp” of a trapezoid pattern (i.e., ramp pattern traced by translating the slider platform upwards with constant velocity). At the new TOP position (reached at the end of every upward translation during trapezoid condition), participants need to hold the slider platform steady for 2s. This would trace the static ‘flat’ portion of the trapezoid pattern. This would then be followed by tracing the “down-ramp” of a trapezoid (i.e., ramp pattern traced by translating the slider platform downwards with constant velocity) that involved translating the thumb back to the HOME position. After reaching the HOME position, the participants had to hold the thumb platform steady at the HOME position for 2s, thereby tracing the static ‘flat’ portion at HOME. In each trial, the participants need to trace three such trapezoid patterns arranged sequentially.

Similarly, in the inverted trapezoid condition, the participants were made to trace the down-ramp of the inverted trapezoid pattern by translating the slider platform 1.5 cm downwards from the HOME position. At the new BOTTOM position (reached at the end of every downward translation during inverted trapezoid condition), participants need to hold the slider platform steady for 2s. This traced the static ‘flat’ portion of the inverted trapezoid pattern. This would then be followed by tracing the up-ramp of the inverted pattern. Thus, it involved translating the thumb back to the HOME position. Again, the participants had to hold the thumb platform steady at the HOME position for 2s, tracing the static ‘flat’ portion at HOME. In each trial, the participants need to trace three such inverted trapezoid patterns arranged sequentially. (see Fig.2B). In both conditions, in each trial, after tracing the last ramp, the participants had to hold the thumb platform steady at HOME position for 3s to complete the trial.

In each condition, there were twelve trials. The duration of each trial was 30 seconds. A minimum rest period of one minute was provided between the trials, and a ten minutes break was provided between the conditions. The order of the conditions was balanced across participants.

### Data analysis

Data analysis was performed offline using Matlab (Version R2016b, MathWorks, USA). Force/Torque data and laser displacement data of thumb were lowpass filtered at 15Hz using second-order, zero phase lag Butterworth filter. There were four static ‘flat’ positions of the thumb: TOP and HOME-TOP during trapezoid condition, HOME-BOTTOM, and BOTTOM during inverted trapezoid condition (refer Fig.2B). In each trial of the two conditions, there were three static ‘flat’ portions for each position. The first one-second force data (of 100 samples) from each of these three static ‘flat’ portions were extracted. Therefore, in total, for a participant, there would be 36 segments (12 trials x 3 segments for each trial) of one-second data for each of the position.

### Root Mean Square Error on the Thumb displacement data

Thumb displacement data collected during both conditions were averaged across trials and participants. We computed the root mean square error (RMSE) for the four static positions (TOP, HOME-TOP, HOME-BOTTOM, BOTTOM) of the thumb displacement data with respect to the template pattern to examine the accuracy of thumb to trace the patterns. RMS error was calculated for each segment, then averaged across 36 segments for each participant, across all participants for each of the four static positions separately.

### Absolute Normal and Tangential force

Normal and tangential forces of the individual fingers and thumb were averaged across 36 segments of all four static positions separately. Then, the normal and tangential force data were averaged across the time samples and participants. The standard error of the mean was also computed.

### Change in the normal forces

The change in the normal force gives information on the difference in the magnitude of forces produced before the start and after the end of each ramp. In each trial of each condition, there were three up-ramps and three down-ramps. In the trapezoid condition, 100 samples of force data immediately before the start of each up-ramp and immediately after the end of that up-ramp were averaged separately. The difference in the mean force was computed. There were 36 such differences (12 trials x 3) for a single participant.

Likewise, the change in the normal forces of all the fingers and thumb were obtained for the down-ramps of trapezoid condition. Similarly, it was computed for both down-ramps and up-ramps of inverted trapezoid conditions separately. Finally, these data were averaged across all participants for various conditions.

### Statistics

We performed all the statistical analysis using R. Two two-way repeated-measures ANOVA was performed on the absolute normal and tangential forces with factors such as static position (levels: TOP, HOME-TOP, HOME-BOTTOM, BOTTOM) and fingers (levels: Index, Middle, Ring, Little). Another two-way repeated-measures ANOVA was performed on the change in the normal force with the factors as movements (levels: UP from HOME and DOWN from HOME) and fingers (levels: Index, Middle, Ring, Little). Sphericity test was done on the data, and the number of degrees of freedom was adjusted by Huynh-Feldt (H-F) criterion wherever required. Pairwise post hoc Tukey tests were performed to examine the significance within factors. Two one-way repeated-measures ANOVA was performed on Thumb normal force and thumb displacement data with factor as static position. Since, by mechanics, the thumb normal force is always dependent on the normal forces of other individual fingers, ANOVA was performed separately for thumb normal force. An equivalence test was performed to check for equivalence between the Thumb normal force at TOP and HOME-TOP static position using the two one-sided t-tests (TOST) approach^20^ for a desired statistical power of 95%. The smallest effect size of interest (SESOI) was chosen as equivalence bounds.

## Results

### Task Performance

Participants were able to trace the trapezoid and inverted trapezoid patterns within the error margin displayed on the template. Throughout the trial, during both conditions, participants maintained the handle in static equilibrium without any visible oscillations in the thumb displacement data. Furthermore, it was observed that the RMS error for the static positions in the inverted trapezoid condition (HOME-BOTTOM: Mean=6.20, SD=0.12; BOTTOM: Mean=18.49, SD=0.33) was significantly (p<0.001) greater than the static positions at the trapezoid condition (HOME-TOP: Mean=5.61, SD=0.24; TOP: Mean=7.14, SD=0.19). Thus, in Figure 3, we could see a comparatively greater standard error of the mean for the thumb displacement data during the inverted trapezoid condition than during the trapezoid condition. The average tilt angles measured at the four static positions was 1.72±0.38°

**Figure 3.**
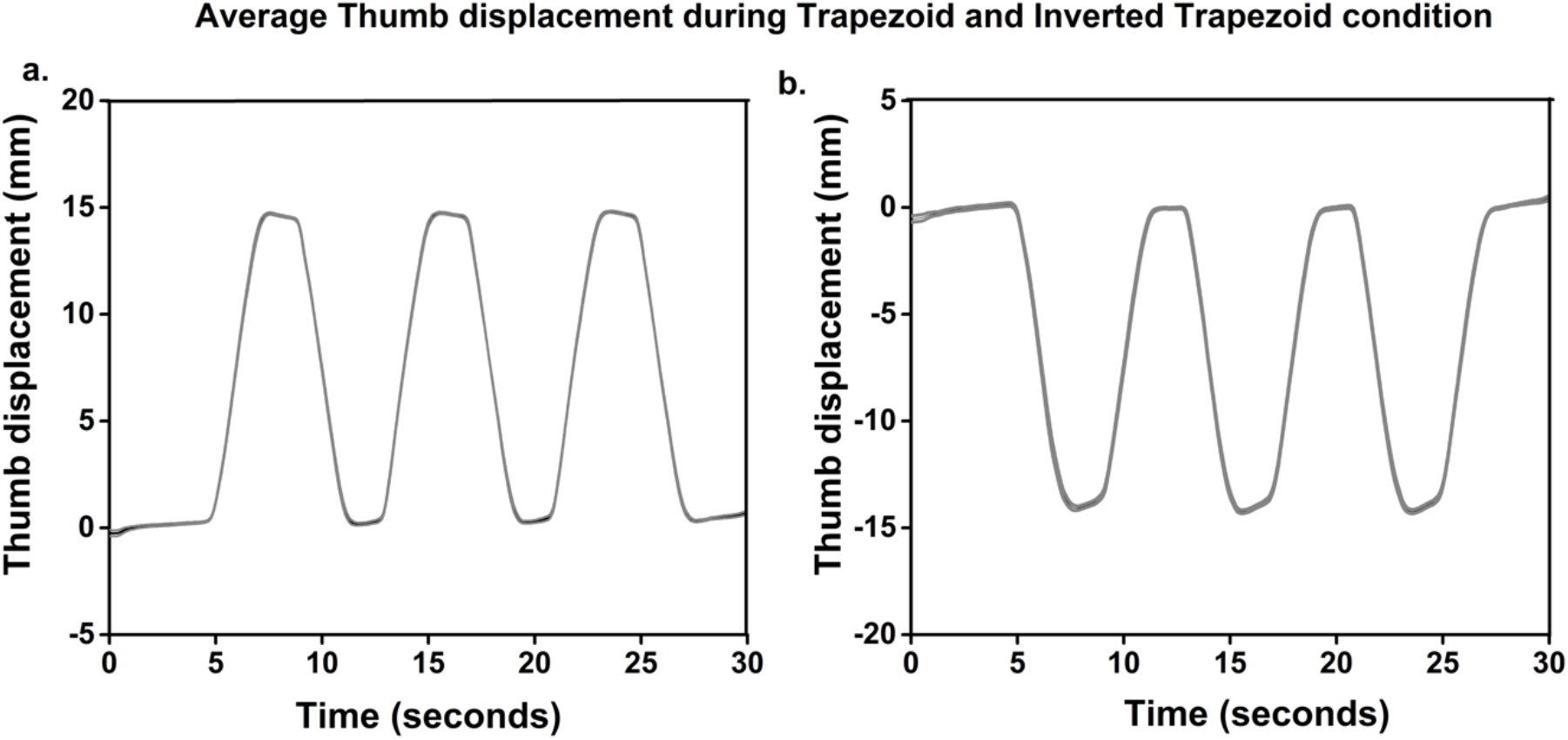
Average time profile of the thumb displacement during trapezoid and inverted trapezoid conditions with the standard error of the mean. The thumb displacement data shown here are averages across trials and participants in each condition. (A) Average thumb displacement during trapezoid condition. (B) Average thumb displacement during inverted trapezoid condition. From the visual observation of the average time profile plots of the inverted trapezoids, it was noticed that the displacement data was not exactly ‘flat’ for the entire two seconds at the BOTTOM position. The first few samples of the BOTTOM portion remained closer to the −1.5 cm, while the remaining samples showed a gradual upward shift of the thumb. However, when the thumb was at TOP, HOME-TOP and HOME-BOTTOM positions, displacement data remained relatively ‘flat’.

### Normal and Tangential forces of fingers and thumb at four static positions

A two-way repeated-measures ANOVA on the normal forces of the individual fingers (except thumb) with the factors fingers and static positions showed a statistically significant effect of fingers (F_(3.63,39.93)_= 40.34; p<0.001, η^2^_p_=0.78) corresponding to a significantly higher (p<0.001) normal force for the little finger followed by the ring finger compared to the radial fingers. There was a significant main effect of the static position (F_(2.61,28.71)_= 32.64; p<0.001, η^2^_p_=0.74) corresponding to a significantly higher normal force (p<0.001) when the thumb was at the BOTTOM position.

The interaction effect of the fingers x static positions was significant (F_(4.27,47.02)_= 187.681; p<0.001, η^2^_p_=0.94), reflecting the fact that the absolute normal force of the little finger when the thumb was at the BOTTOM (7.25N) was significantly greater (p<0.001) than the index finger normal force when the thumb was at the TOP (3.51N) as shown in Fig.4. Among central fingers, the ring finger (3.02N), showed significantly greater (p<0.01) normal force when the thumb remained at the BOTTOM than the middle finger (1.96N) when the thumb reached the TOP.

**Figure 4.**
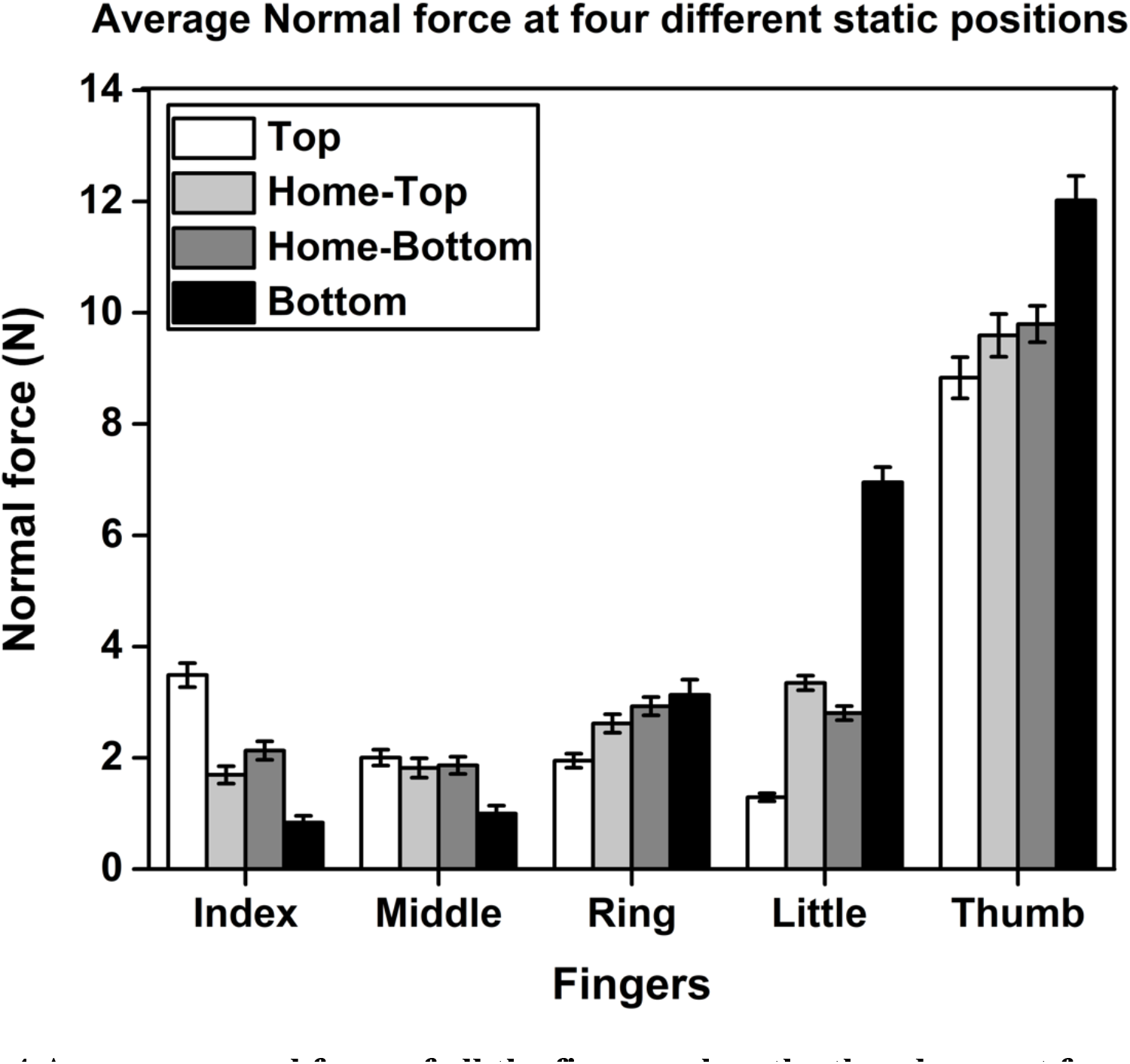
Average normal force of all the fingers when the thumb was at four different static positions. The normal force of the little finger at the static BOTTOM position (black) was significantly greater (p<0.001) than the normal force of the index finger at the static TOP position (white). The index finger exerted significantly greater (p<0.001) normal force than the middle finger when the thumb was at the static TOP position. Likewise, the little finger also exerted significantly greater (p<0.001) normal force than the ring finger when the thumb remained at the static BOTTOM position. Thumb normal force at the static BOTTOM position was significantly greater (p<0.001) than the thumb normal force at static TOP, HOME-TOP (light grey), and HOME-BOTTOM (dark grey) positions.

The normal force of the index finger (3.51N) was significantly higher (p<0.001) than the normal force of the middle finger (1.96N) when the thumb platform was held steady at the TOP. When the thumb reached the HOME position from the TOP during trapezoid condition, index finger normal force (Mean=1.64N, SD=0.54) was not significantly different (t(11) = −1.072, p = 0.307, dz=0.30) from the middle finger normal force (Mean=1.79N, SD=0.57). Thus, the comparison was found to be statistically equivalent (t(11) = 2.530, p = 0.014) as the observed effect size of the dependent means fall within the equivalence bounds of Δ_L_=-1.04 and Δ_U_=1.04.

In the inverted trapezoid condition, while the thumb was tracing the BOTTOM of the inverted trapezoid, little finger normal force (7.25N) was found to be significantly greater (p<0.001) than the ring finger normal force (3.02N). Whereas, when the thumb reached the HOME position from the BOTTOM during inverted trapezoid condition, normal forces of the ulnar fingers were found to be statistically non-significant (t(11) = 0.559, p = 0.588, dz=0.16). The TOST procedure on the dependent pairs (Ring: Mean=2.90N, SD=0.55, Little: Mean=2.78N, SD=0.43) confirmed that they were statistically equivalent with the observed effect size (dz=0.16) that was within the equivalence bounds of Δ_L_=-1.04 and Δ_U_=1.04.

Moreover, pairwise Post hoc Tukey tests confirmed that the little (2.78N) & ring (2.90N) fingers normal forces during inverted trapezoid condition were significantly greater than the middle finger normal force (1.86N, little: p<0.05, ring: p<0.01) when the thumb reached the HOME position from the BOTTOM. Similarly, the ulnar fingers normal forces (while the thumb was at HOME-BOTTOM) were statistically greater than the index (1.64N, ring: p<0.001, little: p<0.001) & middle fingers (1.79N, ring: p<0.01, little: p<0.01) normal forces when the thumb was at HOME-TOP position. Whereas, during the trapezoid condition, when the thumb reached the HOME position, little finger (3.43N) normal force was significantly greater (p<0.001) than the index (1.64N) and middle fingers (1.79N) normal forces.

One-way repeated-measures ANOVA on the thumb normal force showed a significant effect of static position (F_(2.64,29.04)_= 31.90; p<0.001, η^2^_p_=0.74). The thumb normal force at the TOP was not statistically different from the thumb normal force at HOME-TOP. Further, it was confirmed that the thumb normal force at the BOTTOM (12.13N) was significantly (p<0.001) greater than the thumb normal force at the HOME-TOP (9.58N) during trapezoid and HOMEBOTTOM (9.79N) during inverted trapezoid condition.

With regard to the tangential forces, a two way repeated measures ANOVA showed that the factors such as fingers (F_(2.91,32.01)_= 7.50; p<0.001, η^2^_p_=0.40), static positions (F_(2.1,23.1)_= 9.65; p<0.001, η^2^_p_=0.46) and fingers X static positions (F_(4.23,46.53)_= 38.15; p<0.001, η^2^_p_=0.77) interaction affected significantly the tangential forces of the individual fingers (except thumb). When the thumb was at TOP and HOME-TOP during trapezoid condition, the index finger tangential force was neither significantly different nor statistically equivalent to the middle finger tangential force. When the thumb was at the BOTTOM, the little finger tangential force (2.66N) was significantly greater (p<0.001) than the ring finger tangential force (1.13N) (refer Fig.5). Whereas when the thumb returned to the HOME-BOTTOM, ulnar fingers tangential forces (Ring=1.50N, SD=0.37; Little=1.31N, SD=0.49) remained statistically equivalent, with the observed effect size (dz=0.30), lying within the equivalence bounds of Δ_L_=-1.04 and Δ_U_=1.04. At the same position, ulnar fingers tangential forces were found to be significantly greater (p<0.05) than the index finger tangential force (0.62N).

**Figure 5.**
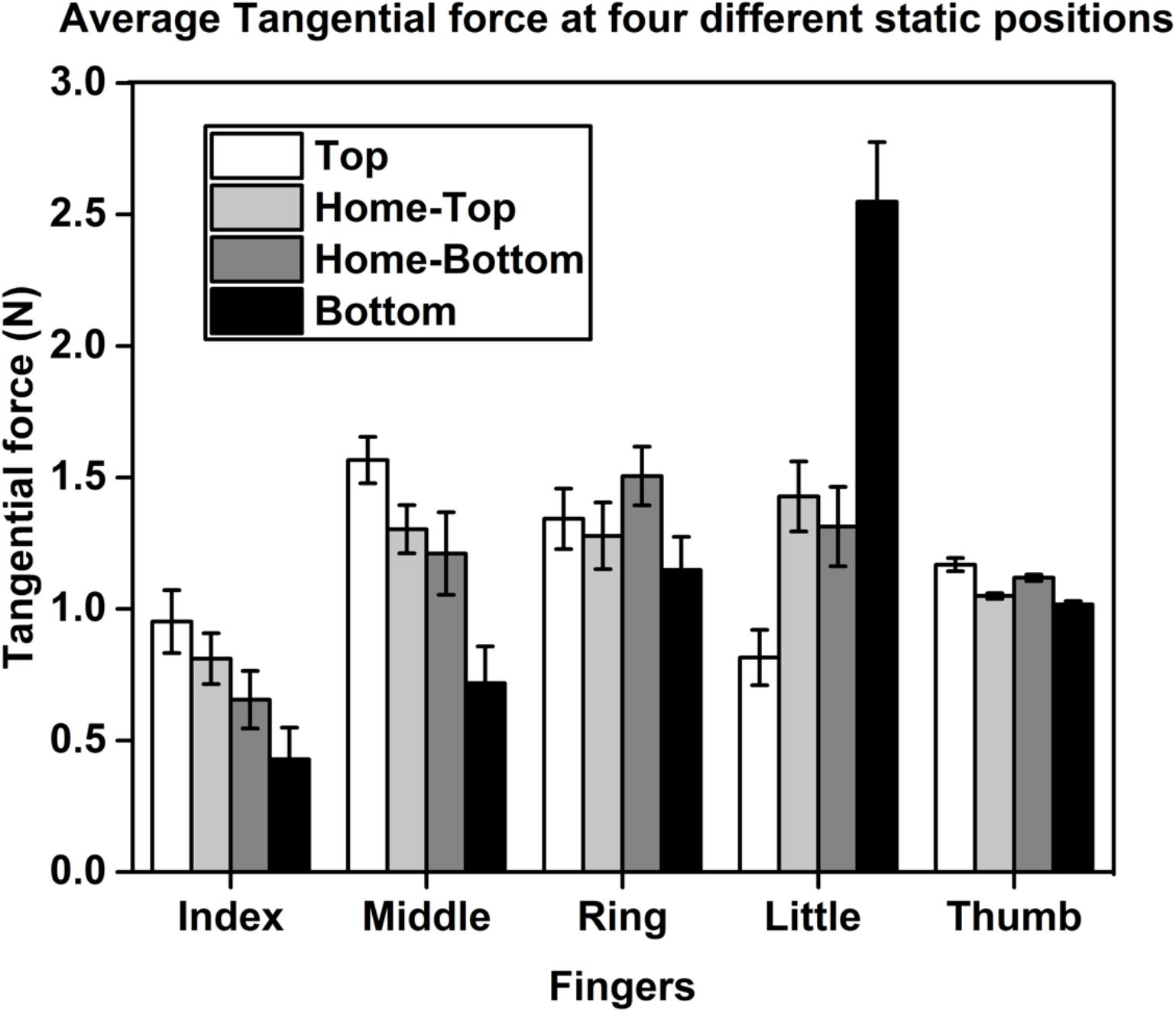
Average tangential force of all the fingers when the thumb was at four different static positions. The little finger tangential force at the static BOTTOM position (black) was found to be significantly greater (p<0.001) than the index finger tangential force when the thumb remained at the static TOP position (white).

Notably, the little finger tangential force (2.66N) (when the thumb was at static BOTTOM position) was significantly greater (p<0.001) than the index finger tangential force (0.92N) (when the thumb was at static TOP position) and both radial fingers (Index: Mean=0.43N, Middle: Mean=0.70N) (when the thumb remained at BOTTOM).

### Change in the normal forces of the individual fingers

From the results of the two-way repeated-measures ANOVA on the change in the normal forces of the individual fingers, we observed a significant main effect of factors such as fingers (F_(2.34, 25.74)_= 78.104; p<0.001, η^2^_p_=0.87), movements (F_(0.83, 9.13)_= 49.83; p<0.001, η^2^_p_=0.81) and their interaction (F_(2.49, 27.39)_= 301.63; p<0.001, η^2^_p_=0.96).

During the upward translation of the thumb from HOME, the change in the normal force of the index finger (1.79N) was significantly different (p<0.001) from the change in the normal forces of the middle (0.008N), ring (−0.88N) and little (−1.84N) fingers (see Fig.6). Likewise, during the downward translation of the thumb from HOME, the change in the normal force of the little finger (4.01N) was significantly greater (p<0.001) than the change in the normal force of the index (−1.07N), middle (−0.93N) and ring (−0.03N) fingers. Since the magnitude of the thumb displacement remained the same in both directions (also, the fingers were equidistant from the center of the handle), the increase in the peripheral fingers normal force was expected to be the same. However, our critical finding was that the change in the normal force of the little finger (4.01N) (when the thumb moved down from HOME) was significantly greater (p<0.001) than the change in the normal force of the index finger (1.79N) (when the thumb moved up from home). Table 1 summarizes the salient results with the ANOVA details.

**Figure 6.**
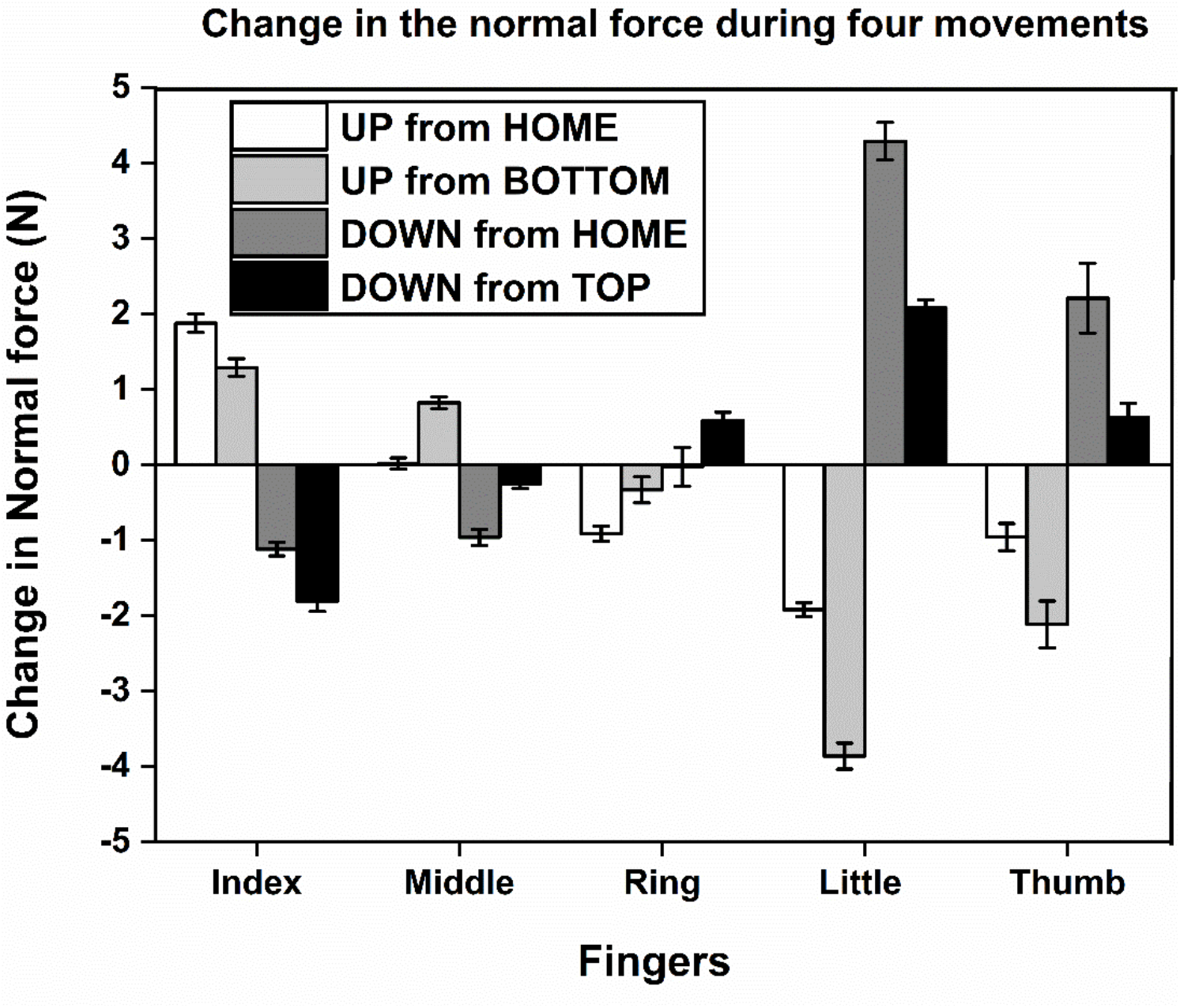
Change in the normal force during four movements of the thumb. The change in the normal force of all fingers and thumb obtained from the up-ramp of the trapezoid condition fall into the category UP from HOME position. In the same way, change in the normal force obtained from the down-ramps of trapezoid condition, up-ramps, and down-ramps of inverted trapezoid condition fall into categories such as DOWN from TOP position, UP from BOTTOM position, and DOWN from HOME position, respectively. The change in the normal force of the little finger during the downward movement of thumb from HOME (DOWN from HOME-dark grey) was significantly greater (p<0.001) than the change in the normal force of the index finger during the upward movement of thumb from HOME (UP from HOME-white).

**Table 1.**
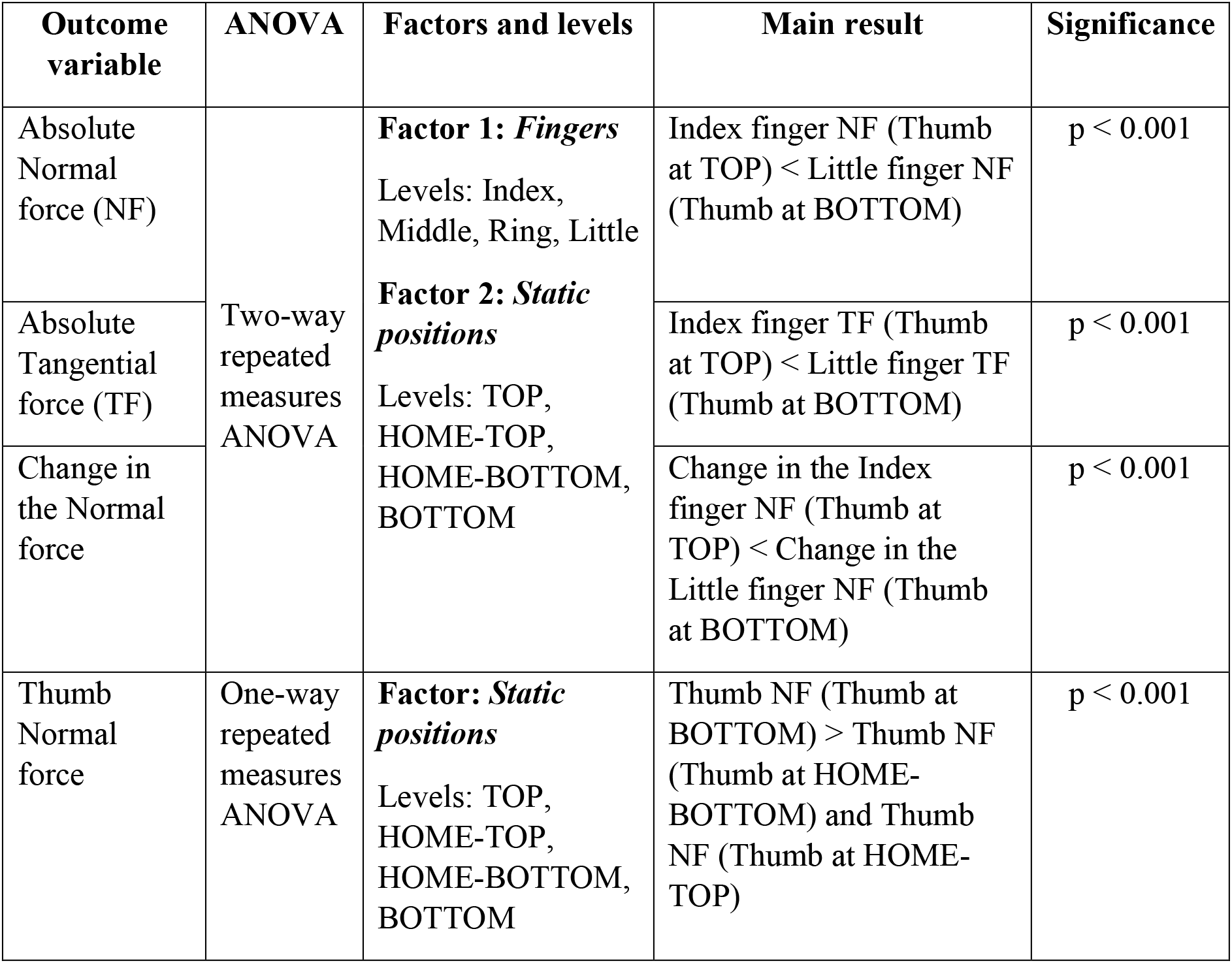
Summary of results with the ANOVA details. The table shows the main result obtained from the statistical analysis (ANOVA) for the outcome variables such as Absolute Normal force, Absolute Tangential force, Change in the Normal force and Thumb Normal force with the significance level.

## Discussion

In this study, we attempted to investigate the contribution of peripheral fingers towards handle stabilization when the thumb platform was vertically translated to different positions. The displacement data of the thumb while tracing the pattern displayed on the monitor served as visual feedback for the participants. The participants were instructed to maintain the static equilibrium of the handle throughout the trial. The displacement of the thumb platform from the HOME position remained the same during trapezoid and inverted trapezoid conditions. Therefore, from mechanics, we posited that the role of the index and little finger for establishing static equilibrium would be similar. However, in contradiction to our expectation, fingertip forces of the peripheral fingers showed different behavior during handle stabilization.

One distinguishing feature of this study was how torque changes were introduced to the grasped handle. Previous studies^21–23^ focused on examining the fingertip forces in handle stabilization when a specific mass was suspended at a certain distance on a horizontal beam attached to the bottom of the handle. This induced external torque changes to the grasped handle, which could ultimately lead to the rotation of the handle. The novelty of the current study is placing the thumb on a slider platform and vertically translating over the railing provided at the handle. The upward and downward translation of the thumb towards and away from the HOME position cause a continuous change in the moment arm of the thumb normal force. Hence, the translation of the movable thumb platform introduced torque changes to the handle. This causes a change to rotational equilibrium of the handle depending on the direction of the thumb displacement.

Also, in previous studies, the external torque changes were of the order of newton-meters (Nm). Therefore, there was a necessity to produce a large compensatory moments to counter-balance. In the current study, the magnitude of thumb displacement was 1.5 cm in either direction. Hence, the magnitude of torque changes was of the order of newton-centimeters (Ncm), which was comparatively lesser than the previous studies. Whether the peripheral fingers behave similarly as in the earlier studies, supporting the MAH during small torque changes is worthy of exploration. Apart from this, the handle utilized in the current study required a movement at the Carpometacarpal (CMC) joint of the thumb for the task execution (i.e., torque changes).

During the trapezoid condition, when the thumb platform reached the TOP position, the index finger produced a greater normal force than the middle finger. Similarly, when the thumb reached the BOTTOM position during the inverted trapezoid condition, the little finger produced a greater normal force than the ring finger. Although the thumb motion caused an increase in the normal force of both the peripheral fingers, we attempted to investigate how these finger forces showed significant variation in detail. As mentioned in the introduction, there were four possible options by which the peripheral fingers normal force pattern may vary (refer to Fig.1).

According to the first and fourth option, the index finger would show a greater change in the normal force when the thumb platform translates from HOME to TOP position than the little finger when the thumb platform translates from HOME to BOTTOM position. In a prehension study^24^ investigating the digit interaction, the percentage share of normal forces contributed by each finger was reported. The results of the study showed that the force shared by the index finger was greater than the little finger, probably because the index finger was the strongest finger. Also, in studies with the handle experiencing external torque changes, the index finger produced greater normal force during pronation moment than the normal force produced by little finger during supination moment^12,22^. As the index finger is stronger than the little finger^25–27^, the *change* in the index finger normal force was expected to be greater than the *change* in the little finger normal force. The only difference is that, in the first option, the absolute normal force of the index finger would be comparable to the little finger. Whereas, in the last option, the absolute normal force of the index finger would be significantly greater than the little finger^10,28^.

Based on the second option, since the thumb displacement is the same in both directions from HOME, the *change* in the normal force of peripheral fingers is expected to be equivalent. However, in reality, the absolute normal force of the peripheral fingers was matching the pattern expected in the third option. According to the third option, the *change* in the normal force and the absolute normal force will be significantly greater for the little finger than the index finger. What could be the reason for the larger increase seen in a weaker finger (little) when compared with a stronger finger (index)? Considering the displacement of the two fingers is the same, this behavior of the little finger is intriguing. We focus on this principal question in the rest of the discussion.

While tracing the up-ramp of the trapezoid pattern, there was an increase in the clockwise moment due to the increase in the moment arm for the normal force of the thumb. Consequently, an anticlockwise moment was produced by increasing the normal force of the radial fingers. In an earlier study^7^, the force sharing pattern for normal forces of the first four fingers varied while the location of the thumb was changed discretely. It was found that the normal force of the index and middle finger increased when the thumb was positioned opposite to the middle finger. Our results are in broad agreement with these results. Since the ring and little fingers produce a clockwise moment, the normal force of the ring (around 1N) and little finger (around 2N) decreased during the up-ramp of the trapezoid.

Furthermore, the tangential force of the thumb increased slightly (around 0.10N) during the upward translation of the thumb from the HOME position due to the contribution of inertial force along with the gravitational force^29,30^. The increase in the tangential force of the thumb has to be compensated by a corresponding decrease in the tangential force of the virtual finger (VF). Among the VF, the drop in the tangential force could have been evenly shared within all the fingers or with ulnar fingers alone. Instead, a notable drop in the tangential force was seen only in the little finger. The reason for the little finger to decrease its tangential force during the upward translation of the thumb may be explained from a biomechanics perspective. The upward movement of the thumb while holding an object is considered as an extension movement or radial abduction of the CMC joint of the thumb. This movement happens due to the contraction of muscles such as abductor pollicis longus and extensor pollicis brevis^31^.

Abductor pollicis longus, a primary radial deviator of the wrist, contracts causing radial deviation of the wrist joint^32^. Hence, we believe that the radial deviation caused by the radial abduction of the thumb could be resisted by a possible contraction of the abductor digiti minimi of the little finger as it could cause ulnar deviation of the wrist joint^33^. In addition to the anatomical restriction of the wrist motion, externally, the participant’s wrist was strapped by using velcro to arrest any unwanted movement of the wrist. Thus, we believe that the radial abduction of the thumb while grasping a handle may have resulted in the little finger abduction in the form of medial rotation. Abduction of the little finger was indirectly noticeable from the decrease in the little finger tangential force. Consequently, tangential and normal forces of the index, middle, and ring fingers increased slightly.

During the downward translation of the thumb during both conditions, the tangential force of the thumb decreased slightly by about 0.2N. Subsequently, the tangential force of the other fingers increased to balance the vertical equilibrium. From the mechanics standpoint, if there were an increase in the tangential force of the radial fingers, it would be accompanied by the increase in the normal force of the same finger to prevent slip^34^. Thus, it would cause a tilt in the anticlockwise direction, adding up to the tilt caused due to the downward shift of the thumb to the BOTTOM position. The increase in the tangential force could have been shared by the ulnar fingers. But, in reality, there was a significant increase in the tangential force of only the little finger, while the ring finger showed a drop in the tangential force. This was contrary to our expectations. Since the change in the forces exerted by the little finger was quite prominent, the forces exerted by the other fingers reduced.

The downward translation of the thumb towards the little finger may be considered a full flexion or opposition movement^35^. Opponens pollicis of the thumb is responsible for this movement^36^. It contracts during the downward translation of the thumb to BOTTOM position. Meanwhile, opponens digiti minimi of the little finger acts in synergy with the opponens pollicis longus^37^. It is known that the opponens digiti minimi is one of the antagonist muscles of the opponens pollicis^38^. Thus, there is a possibility for the little finger to produce lateral rotation^36^, which occurs in the form of upward displacement of the point of force application (or finger ‘rolling’ within the sensor) of the little finger towards the ring finger. This could have caused an increase in the little finger’s tangential force while the other fingers tangential force dropped to maintain the vertical equilibrium.

Subsequently, the increase in the tangential force of the little finger would be accompanied by an increase in the normal force of the little finger. Thus, the full flexion (downward translation of the thumb from HOME position) of the thumb’s CMC joint may have caused simultaneous adduction of the little finger (i.e., the movement of the little finger towards the ring finger side). This is indirectly apparent from the tangential forces. Conversely, such a rise in the index finger tangential force was not seen when the thumb translated to the TOP position. The radial abduction of the thumb did not cause a considerable amount of abduction of the index finger, as both the normal and tangential forces of the adjacent middle finger contributed towards exerting a pronation moment. So, the involvement of the index finger tangential force was slightly less in the total tangential force.

Thus, the downward movement of the thumb caused a substantial increase in the adduction of the little finger, probably due to morphological reasons. We found a significantly greater normal force produced by the little finger during the supination moment than the normal force exerted by the index finger during the pronation moment. Consequently, due to the higher increase in the little finger normal force, thumb normal force increased to 12N during the downward translation to compensate for the clockwise tilt caused. In the case of upward translation of thumb from HOME to the TOP position, thumb normal force was 8N because the change in the index finger normal force was not high enough. Thus, the little finger forces are different from index finger forces during the thumb movement towards them. Further investigation on the anatomical relationship between the thumb and little finger is necessary to better understand this distinct behavior of the little finger.

## Concluding comments

When an unsteady thumb platform translated to different positions vertically, the normal forces exerted by the individual fingers varied in a systematic way. In particular, if the thumb platform remained static at various positions, it caused remarkable changes in the forces of peripheral fingers for object stabilization. The forces and change in the force produced by the little finger when the thumb was at the BOTTOM position were comparatively greater than those of the index finger when the thumb was at the TOP position. This distinct behavior of the little finger compared to the other fingers perhaps suggests that there is an anatomical/morphological relationship between the thumb and little finger. Further investigation on the anatomical function during the task performance may give a better idea of this relationship.

## Acknowledgments

We thank the Department of Science & Technology, Government of India, for supporting this work, vide Reference Nos SR/CSRI/97/2014 & DST/CSRI/2017/87 under Cognitive Science Research Initiative (CSRI) (awarded to Varadhan SKM) and American Express (funding awarded to DART lab, IIT Madras). The funders had no role in study design, data collection and analysis, decision to publish, or preparation of the manuscript.

## Author Contributions Statement

Conceptualization – VSKM; Methodology – BR, VSKM; Formal Analysis – BR; Writing Original Draft – BR; Writing, Review, and Editing – BR, VSKM; Funding acquisition – VSKM.

## Data Availability

We plan to publish a data descriptor article along with this manuscript. Hence the data will be made available in due course of time.

